# GLMsim: a GLM-based single cell RNA-seq simulator incorporating batch and biological effects

**DOI:** 10.1101/2024.03.20.586030

**Authors:** Jianan Wang, Lizhong Chen, Rachel Thijssen, Belinda Phipson, Terence P. Speed

**Affiliations:** Bioinformatics Division, Walter and Eliza Hall Institute of Medical Research, 1G Royal Parade, Melbourne, 3052, VIC, Australia; School of Mathematics and Statistics, University of Melbourne, Grattan Street, Melbourne, 3010, VIC, Australia; Blood Cells and Blood Cancer Division, Walter and Eliza Hall Institute of Medical Research, 1G Royal Parade, Melbourne, 3052, VIC, Australia

**Keywords:** scRNA-seq, simulation, generalized linear model, library size, batch, biology, associations

## Abstract

With development of the single cell RNA-seq technologies, large numbers of cells can now be routinely sequenced by different platforms. This requires us to choose an efficient integration tool to merge those cells, and computational simulators to help benchmark and assess the performance of these tools. Although existing single cell RNA-seq simulators can simulate library size, biological and batch effects separately, they currently do not capture associations among these three factors. Here we present GLMsim, the first single cell RNA-seq simulator to simultaneously capture the library size, biology and unwanted variation and their associations via a generalized linear model, and to simulate data resembling the original experimental data in these respects. GLMsim is capable of quantitatively benchmarking different single cell integration methods, and assessing their abilities to retain biology and remove library size and batch effects.

## Introduction

During the past few years, there has been significant progress in single cell sequencing technologies, which allow scientists to explore gene expression at individual cell resolution. In contrast to bulk-RNA seq that studies average expression at the cell population level, single cell RNA-seq technologies are capable of exploring the transcriptomics of each individual cell. Advanced single cell protocols offer researchers new ways to understand biologically and medically relevant questions, such as the response of immune cells to anti-tumour drugs[1], the dynamic progress of embryonic cell state evolution[2], cellular compositional changes across healthy and diseased tissues[3], detection of pathogenic pathways for neurodegenerative patients[4], and the potential regulatory gene changes of diabetes patients[5].

As single cell transcript sequencing became more popular, a rapid growth of tools occurred to answer different biological questions. The online single cell RNA sequencing (scRNA-seq) tool database[6] has recorded more than 1440 software packages available for different analysis tasks. The choice of the tools determines the analysis results; therefore, using the most appropriate tool to carry out an analysis is essential for researchers. Benchmarking, which applies a number of tools to several datasets to decide the best-performing tool, is a direct solution to this multiple-choice issue. Despite remarkable progress having been made, benchmarking studies still face great challenges. One of the challenges is to find suitable datasets, because benchmarking results are highly dependent on the datasets that are used[7]. Inappropriate datasets can lead to biased selection of tools. Hence, researchers should make sure that each dataset is suitable for evaluating different methods. Another challenge is relevant to the characteristics of single cell data[8]. The complex structure of single cell data makes it hard to get complete information from experimental data directly. Unknown information left in the dataset can undermine the objectivity of the benchmarking results.

Simulation in this context refers to the creation of a computational model which represents and displays essential characteristics of real-world single cell RNA-seq data. Access to a faithful simulated dataset is of vital importance for the conduct of comparative studies and to help developers to check their methods[9–13]. Such assessments are difficult to carry out on the original single cell data if the experiments are not specifically designed. Simulation, however, can easily realize different scenarios and create extreme cases that can be used for testing. Simulation provides developers with an opportunity to investigate the influence of different parameters, examine the robustness of a method and validate the assumptions behind the method. In addition, it is hard to obtain real data with a wide range of conditions due to limited budgets. Simulation is much less costly than real biological experiments. Lastly, the quality of the original data is always doubtful, such as when the experiment damages droplets and lyses cells. Such low-quality real data will influence the accuracy of benchmarking results. However, simulation is able to address this issue via quality control at the beginning and produce high quality data.

Existing single cell simulation strategies fall into two major categories. One is based on the distributional models. Some popular methods, such as Splatter[14], SPARSim[15], SPsimSeq[16], scDesign[17], scDesign2[18], POWSC[19] and powsimR[20] use this strategy. These methods fit the data into some statistical distributions, obtain the parameter estimates, and then sample random values from the fitted distributions. Even though this strategy is quick and easy to follow step by step, it is not always known how well the estimated parameters fit the data. If the data fail to fit the distributions, simulation using the parameter estimates will lead to synthetic data which diverges from the original data. The second type of simulator, such as SymSim[21] and Minnow[22], models the key steps in RNA synthesis and in the sequencing process such as the enrichment of transcripts, the polymerase chain reaction (PCR), and molecular fragmentation. This type of model succeeds in quantifying technical errors from the beginning of the sequencing procedure, yet still finds it difficult to accurately simulate all steps of gene expression.

One of the essential applications for the simulated data is to benchmark different single cell data integration methods. Single cell RNA sequencing has been widely applied during the past decade for its strength in exploring biology at single cell resolution. Plenty of integration tools[23–28] designed for single cell analysis purposes have been developed, but none of above-mentioned tools use simulated data to examine their method. As a result, those methods require reliable simulated datasets to evaluate their performances. That is because the unknown ground truth makes it hard to use the original data to finish the assessment task and provide a clear explanation of benchmarking results. In addition, benchmarking different integration methods calls for consideration of the library size, batch and biological information and their associations, because a good integration method is expected to keep the biological differences and remove library size effects and the batch differences. Unfortunately, existing single cell simulators do not satisfy all those conditions. Hence, synthetic data with suitably designed information is necessary to help evaluate the performance of different integration tools.

Although most existing single cell simulation methods[14–16, 18, 21] satisfactorily simulate library size, biology and batches separately, none of the methods currently simulate the associations among the three factors. Here, we present GLMsim (**G**eneralized **L**inear **M**odel based **sim**ulator), a single cell simulator which aims to tackle this issue. GLMsim fits each gene’s counts into a negative binomial generalized linear model (GLM), estimating mean gene expression as a function of the estimated library size, biology and batch parameters and then samples counts from negative binomial distributions. If outlier values arise in the initial simulation, GLMsim has procedures to set outliers back to a standard level. Overall, GLMsim outperforms other methods in producing single cells counts which resembling those in the original data, as a GLM is robust in capturing most essential characteristics of single cell data.

## Results

### An overview of GLMsim framework

Briefly, GLMsim includes three steps: (1) estimating parameters for each gene, (2) simulating single cell gene counts and (3) rescuing outlier genes (Fig. 1). Initially, GLMsim starts from an observed scRNA-seq count matrix that includes the cell type and batch information. GLMsim captures the main characteristics of the data by fitting a generalized linear model, returning estimated parameter values for each gene. Finally, a synthetic count matrix with same number of genes and cells is generated using the estimated coefficients from the previous step. In order to retain most of the essential properties from the original data, GLMsim simulates gene by gene and keeps the same library size as that data. Since it is possible to get outlier values from the simulation, an additional step in GLMsim checks and corrects for outliers if they exist after the initial simulation.

**Fig. 1.**
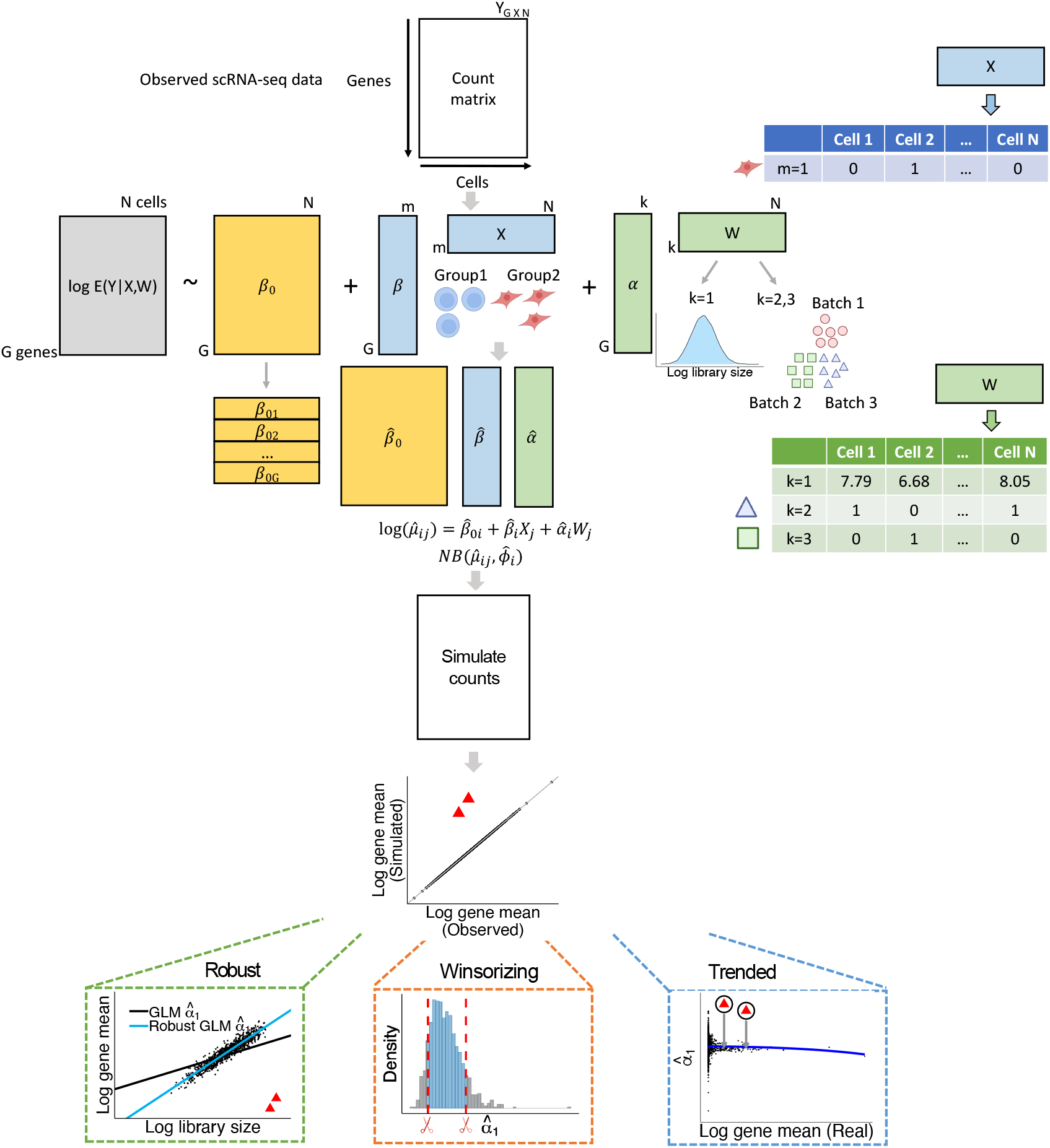
Overview of GLMsim. GLMsim starts from an observed scRNA-seq count matrix, where rows are genes and columns are cells. For each gene, GLMsim applies a generalized linear model to estimate biological and technical coefficients. Next, GLMsim samples single cell counts from a negative binomial distribution with a mean computed using previously estimated coefficients, and dispersion estimated from the observed data. In the last optional step, GLMsim checks if outlier genes exist, and uses one of the three alternative methods to deal with outliers: robust negative binomial GLM, winsorizing the coefficients and trending the coefficients.

### GLMsim captures associations between library size, biological effects and batch effects

In order to evaluate the performance of single cell integration methods, it is important to simulate scRNA-seq data that captures library size, biology and batches. Library size refers to the sum across genes of all counts within a cell. Biology commonly stands for cell types, subtypes or conditions, such as control or stimulated states. Batch factors here are technical variations that cause differences in gene expression across cells, such as different storage conditions, lab operations, protocols or sequencing platforms. Current simulation methods capture biology, some capture batches, while others are good at simulating library size. None of the methods seem to be able to simulate the associations between these three factors (Table 1). Association here means that one factor has potentially different effect across different categories of another factor. For example, cells from different biological groups may have different library size distributions, which is an association between library size and biology. It is also likely that each batch includes heterogenous cell types. Similarly, one cell type may belong to multiple but not all batches. These two scenarios are regarded as involving an association between batch and biology. If the cells with two out of three factors satisfy any of the above conditions, we consider that these associations potentially exist in the dataset.

**Table 1.** Summary of some single cell simulators.

| Method | Library | Batch | Biology | Library × Batch | Library × Biology | Batch × Biology |
| --- | --- | --- | --- | --- | --- | --- |
| Splatter | ✓ | ✓ | ✓ | × | × | × |
| SymSim | × | ✓ | ✓ | × | × | × |
| SPARSim | ✓ | ✓ | ✓ | × | × | × |
| scDesign2 | × | × | ✓ | × | × | × |

Library: library size. Library × Batch: library size associated with batch. Library × Biology: library size associated with biology. Batch × Biology: batch associated with biology.

The Celline2 dataset is a typical one showing association between batch and biology. In the original dataset, batch 3 includes cells from both the Jurkat and 293T cell lines (Fig. 2a). GLMsim enables one to retain the state of mixture of batch 2 and batch 3 cells within the 293T group (Fig. 2b), while Splatter divides the cells into 2 groups and 3 batches without showing the partial mixture between batch 2 and batch 3 cells (Fig. 2c). scDesign2 can simulate the two biological groups, but the isolated pattern between Jurkat cells is not shown (Fig. 2d). SymSim embedded three batch groups into each biological group, and the three batch groups do not have any overlap with each other (Fig. 2e). SPARSim did not represent the batch effect as seen in the original data, though it did demonstrate an association between batch and biology in batch 3 group (Fig. 2f).

**Fig. 2.**
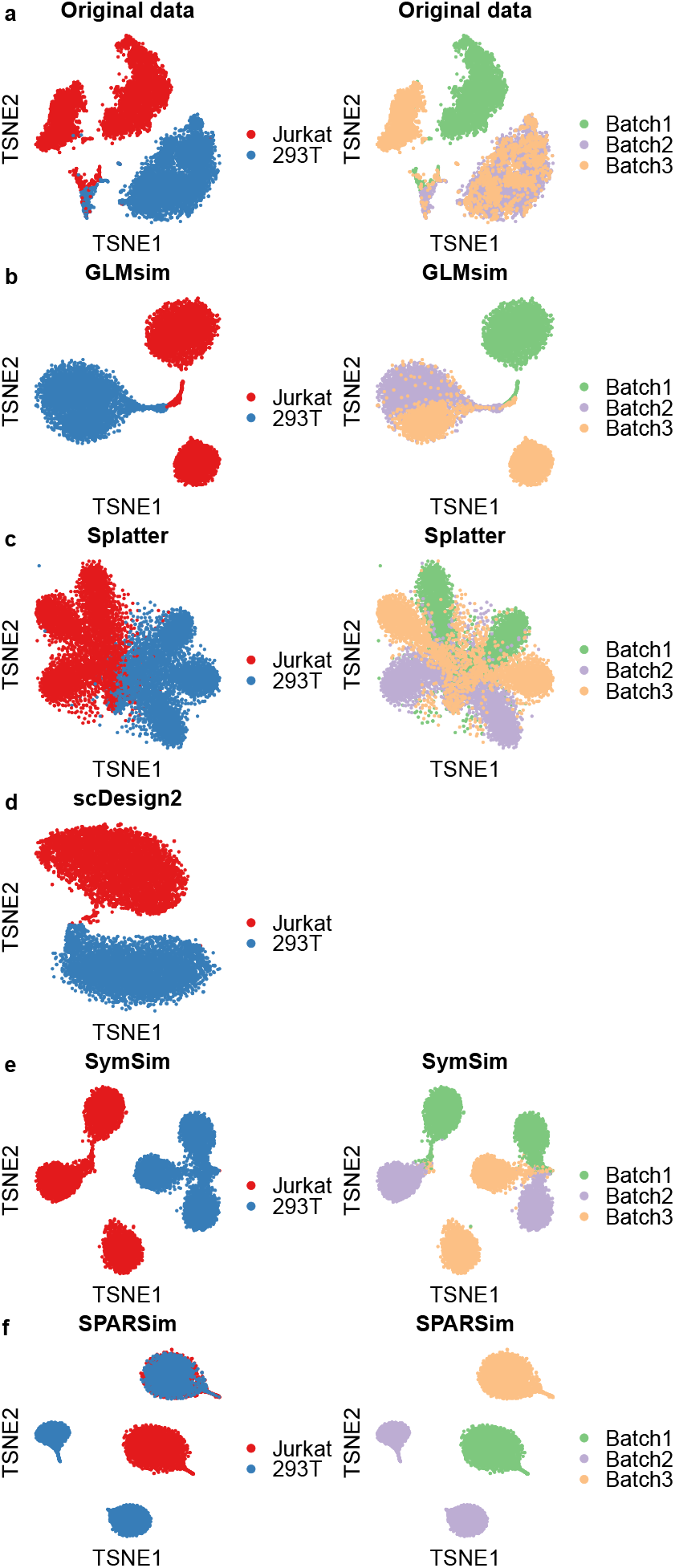
The cellline2 experimental data and data simulated by different methods. Association exists between biological and batch effects. (a-f) tSNE plot of real and simulated data colored by biological groups and batch groups. (a) Real data. (b) GLMsim simulated data. (c) Splatter simulated data. (d) scDesign2 simulated data. Since scDesign2 cannot simulate batch effect, the tSNE plot colored by batch is not shown here. (e) SymSim simulated data. (f) SPARSim simulated data.

In order to explore associations between all three factors, we examine the CLL dataset. It involves 6 biological groups and batches originating from 7 CEL-Seq2 plates. The key point in this dataset is that associations exist between all pairs of the three factors (Fig. 3a,b). None of the popular single cell simulators can deal with this complex dataset. GLMsim simulates data that most resembles the original data (Fig. 3c). For the Granta biological group, GLMsim simulates cells from LC89, LC91, LC93, LC95, LC96, and LC99. Further, a cluster of cells are surrounded by VEN and other treatments, and GLMsim shows a mixture of the groups of cells. Splatter completely failed to simulate this dataset accurately (Fig. 3d), because cells from all batches and all groups were mixed together. Due to the absence of batch effects in their model, scDesign2 can only capture the biology. Gaussian copulas permit scDesign2 to capture part of the biology well, such as in the DMSO and Granta groups, but it is not able to distinguish certain subgroup differences, such as the VEN and VEN+BCLxLi groups that were separated from VEN+MCL1i and VEN+NAV in the original data (Fig. 3e). The performance of SymSim is strange. It splits batches into equal size groups 5,6 and 7, and evenly divided groups 2, 3 and 4 into all batches (Fig. 3f). In addition, the majority of cells have high library sizes which suggests that SymSim failed to simulate the library size effect. The performance of SPARSim is next best after GLMsim (Fig. 3g). SPARSim is able to partition cells into the different biological and batch groups, but it mixed cells in the wrong way. For example, the Granta cells are separated from DMSO cells in the original data, but were mixed together by SPARSim.

**Fig. 3.**
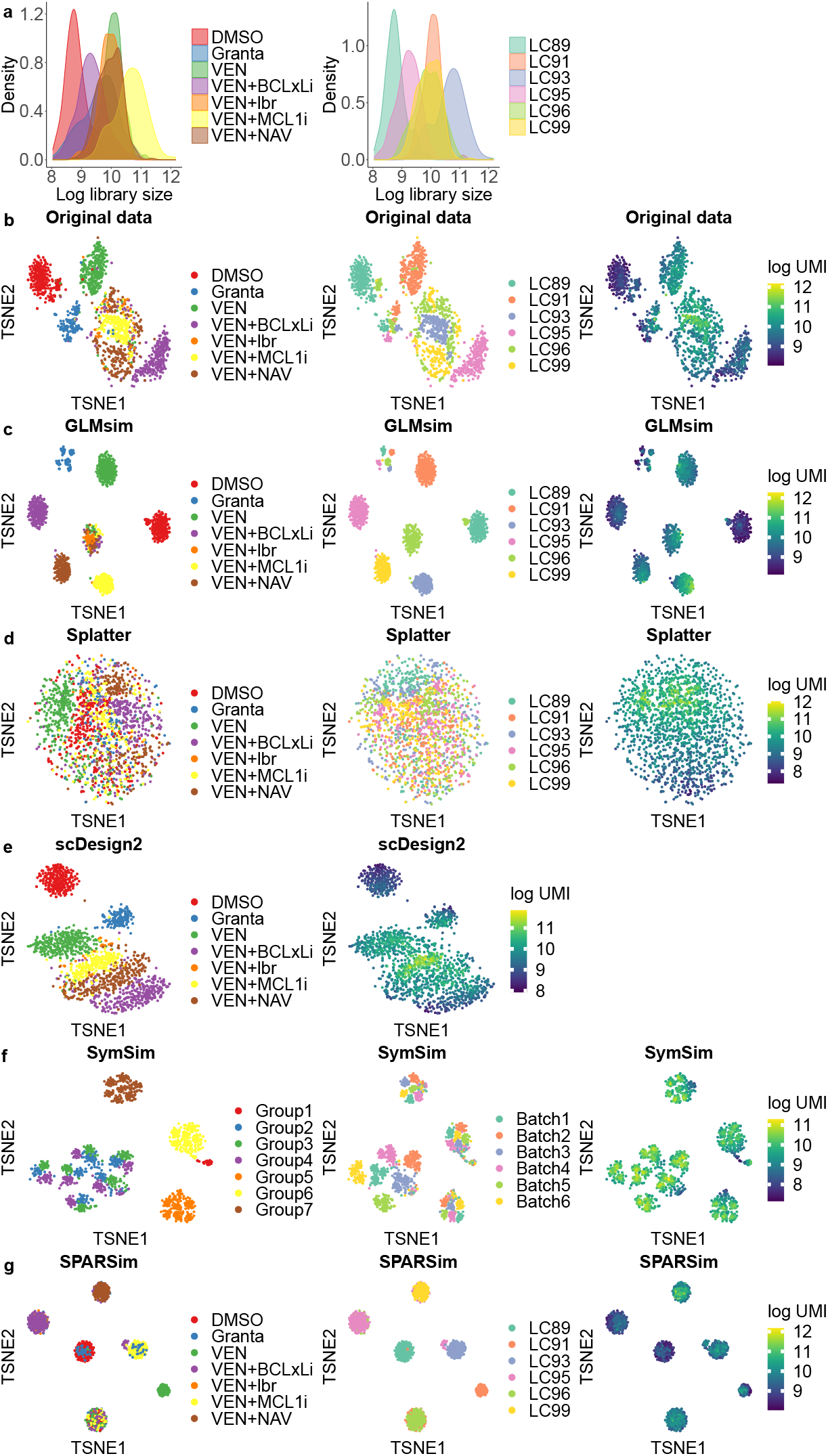
The CLL experimental data and data simulated by different methods. Associations exist between library size, biological and batch effects. (a) The log library size distribution across different biological groups and batch groups. (b-g) tSNE plots of real and simulated data colored by biological groups, batch groups and library sizes. (b) Real data. (c) GLMsim simulated data. (d) Splatter simulated data. (e) scDesign2 simulated data. Since scDesign2 cannot simulate the batch effect, the tSNE plot colored by batch is not shown here. (f) SymSim simulated data. (g) SPARSim simulated data.

The success of GLMsim in simulating associations between library size, biological effects and batch effects using GLMsim lies in its ability to accurately capture biological and unwanted variation simultaneously from original data. The failure of other methods to do so is mainly caused by their estimating library size, biological and batch parameters separately. For instance, Splatter simulates these factors in three independent steps. The implicit assumption of Splatter is that the three factors are independent, and hence it cannot capture associations among the three factors. Further, sampling DE genes for biological and batch factors together will lead to overlap of DE genes among the biological or batch groups and disordered gene rankings for the two factors, and this leads to its inability to simulate complex datasets with multiple biological groups and batches. Gaussian copulas are a powerful strategy to capture gene-gene correlations, and so scDesign2 is able to simulate heterogenous cell clusters. However, the absence of batch terms in the model implies that scDesign2 is unable to simulate a dataset with technical variation. SymSim cannot mimic any given data set for two reasons: (1) It estimates parameters from its own database instead of the given dataset, which limits its simulation ability; (2) It simulates biological and batch effects in sequential order, which prevents it considering the effects jointly. The performance of SPARSim is better, because it simulates library size, mean gene expression and biological variability independently for each cell type. While the SPARSim simulation is competitive, it still includes different groups of cells not present in the original data. That is probably caused by its assumption of a common distribution for batch effects, even if different batches display different features.

### GLMsim keeps most of the essential features of real data

We now compare the simulated data from GLMsim and 4 other single cell RNA-seq simulators on all 6 reference datasets. Utilizing gene-level and cell-level metrics (see “Methods”), we systematically evaluate the performance of all 5 simulators. The gene-level metrics consist of gene means, total gene UMIs, gene variances and gene proportions of zeros, and the cell-wise metrics include library size and cell proportions of zeros.

The gene-level comparisons are all scatter plots between features of the simulated and the reference data. We see that GLMsim outperforms the other methods (Fig. 4a-d). The similarity between GLMsim and the reference data indicates that GLMsim maintains the gene-wise features. scDesign2 and SPARSim ranked in the middle in these comparisons, with small deviations from the reference datasets for several genes (Fig. 4b,c), suggesting that the gene-gene correlations of scDesign2 and individual gamma distributions pf SPARSim can effectively recapitulate gene-level properties. Splatter and SymSim consistently have poor performances for all comparisons and all datasets (Fig. 4a-d). This is due to the fact that Splatter simulates mean gene expression across all genes via a common gamma distribution, while SymSim simulates counts by mimicking the mRNA capturing procedure. In addition, we also investigate how well the methods preserve the relationship between gene means and gene variances as well as gene proportions of zeros (Fig. S1). The non-UMI hESC data has a wider range of mean-variance and mean-zero proportions compared to the UMI datasets.

**Fig. 4.**
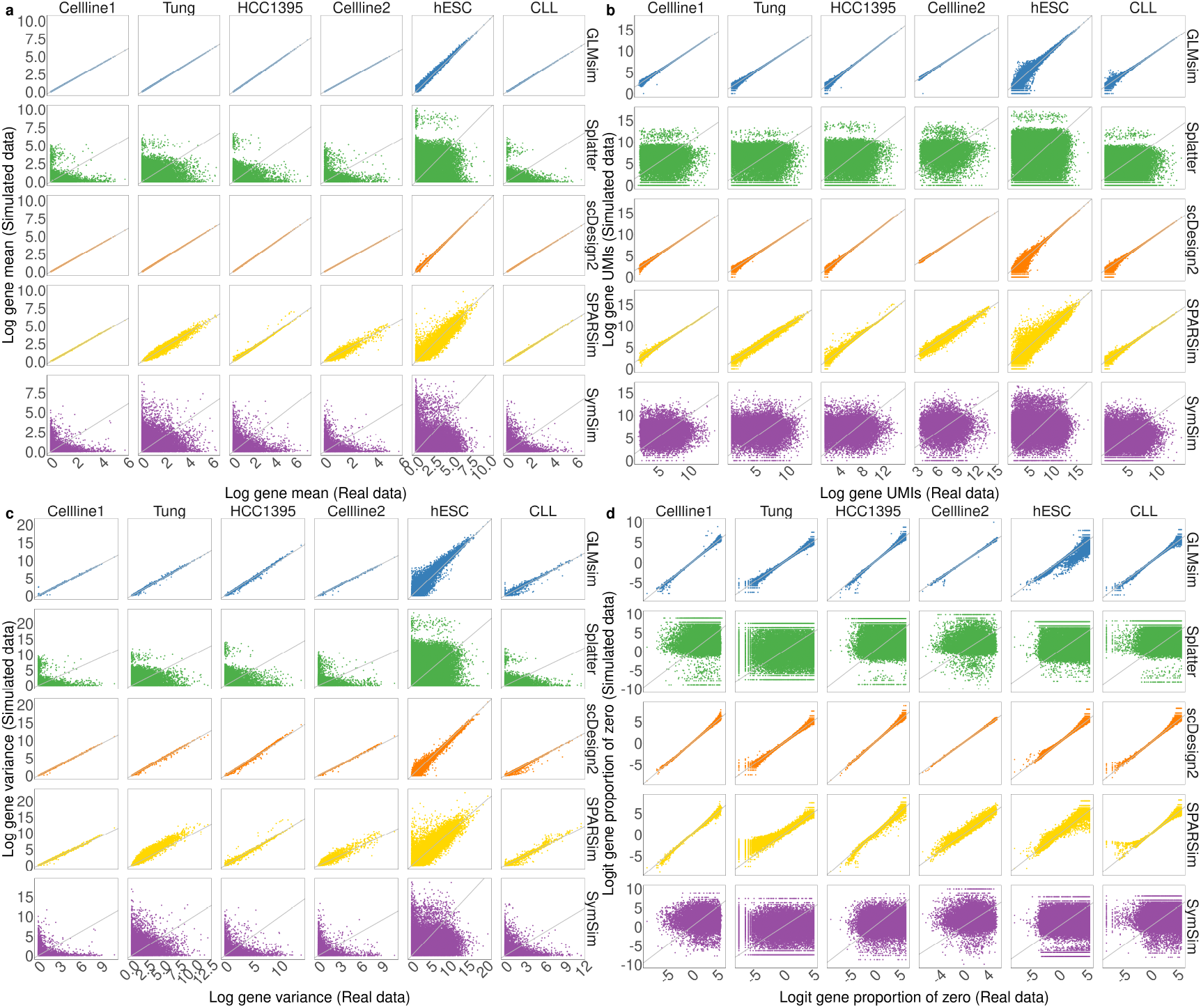
Pairwise comparison of gene-specific features between simulated data and original data. Each row represents a simulation method. Each column represents a dataset. The *x* axis of each plot refers to the metric from the original data, and the *y* axis refers to the metric computed from the simulated data. Each dot represents a gene. (a) Log gene mean. (b) Log gene UMIs. (c) Log gene variance. (d) Logit gene proportion of zero.

For the cell-level comparisons (Fig. 5a,b), GLMsim ranked best, and SPARSim ranked second in simulation of library sizes and cell proportions of zero. SPARSim performed well in simulation of simple data, such as Celline1, Tung, and HCC1395 datasets, but it failed to simulate more complex datasets, which indicates that sampling DE factors from a common distribution limits its ability to simulate complicated situations, despite the fact that the multivariate hypergeometric distribution can handle simple cases. On the contrary, GLMsim incorporates library sizes from reference data directly in the model and recovers cell-level properties. Moreover, GLMsim is the only method successfully simulating the cell proportions of zero on the non-UMI hESC dataset, whereas all other methods show a large difference from this reference data. The remaining methods show weakness in keeping the cell-level attributes. For Splatter, the log normal distribution does not preserve the original library size information, and the logistic regression has drawbacks shaping the dropout events. For scDesign2 and SymSim, their failure is not unexpected since they do not consider library size in their model.

**Fig. 5.**
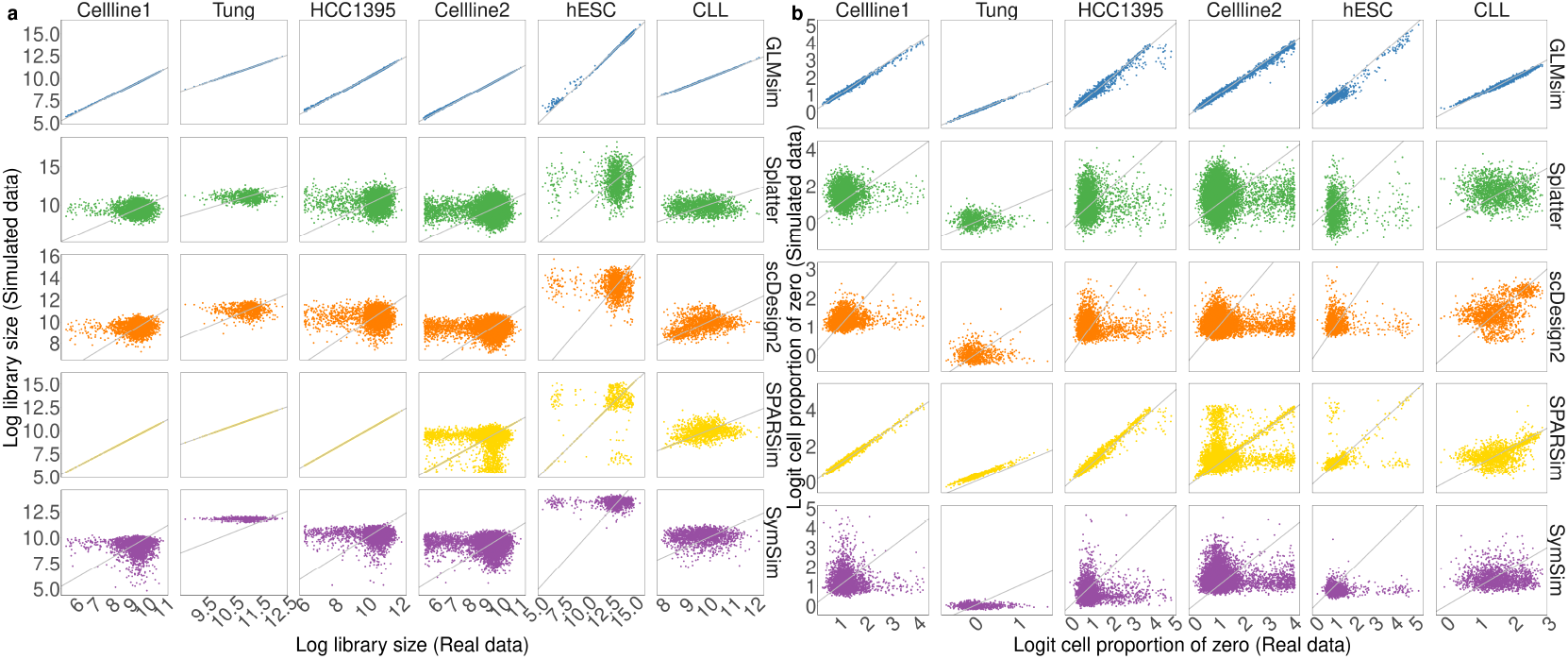
Pairwise comparison of cell-wise metrics between simulated data and original data. Each row represents a simulation method. Each column represents a dataset. The *x* axis of each plot refers to the metric from the original data, and the *y* axis refers to the metric computing from the simulated data. Each dot represents a cell. (a) Log library size. (b) Logit cell proportion of zero.

Overall, GLMsim is better than the other methods in simulating data similar to the reference data in the features we present for both UMI and non-UMI dataset (Fig. 6). SPARSim and scDesign2 perform well on gene-level metrics but fail to capture certain cell-level characteristics, such as library sizes and cell proportions of zero. Splatter and SymSim consistently have lower Spearman correlation with the real data, indicating that the two methods are unable to reproduce data similar to the reference data. The poor performance of these two methods suggests that sampling single cell characteristics from statistical distributions will distort the shape of the simulated datasets as a whole. Additionally, the failure of SymSim lies in estimating parameters from its internal database with the dataset that most similar to the real data, instead of obtaining the parameters from the reference data directly.

**Fig. 6.**
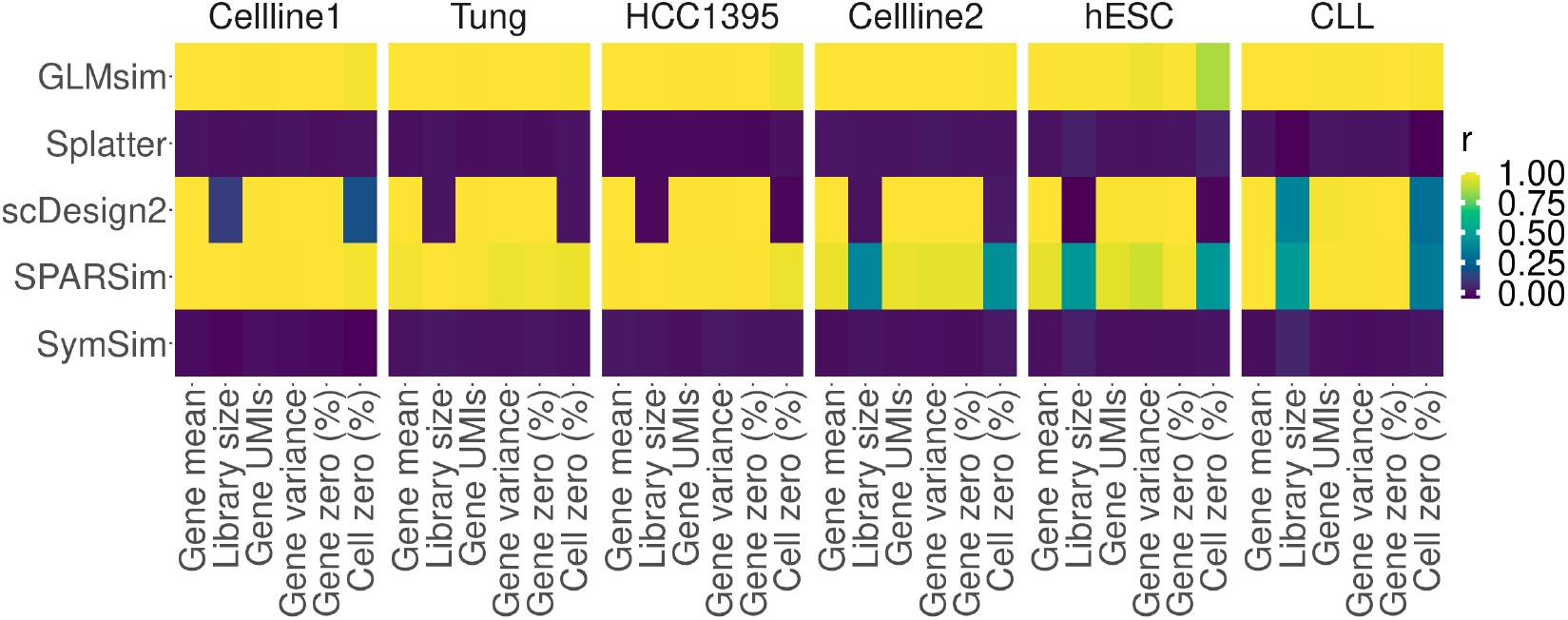
Spearman correlations between features of the simulated data and reference data. Each column stands for a gene- or cell-level metric. Each row stands for a simulation method. The column panels display different datasets. The heatmap is colored by Spearman correlation of a metric between the simulated data and the reference data.

### GLMsim has high computational efficiency

The computational scalability varied across different datasets (Fig. 7). Most of the methods finish the simulation tasks using less than 3 hours CPU time and 10 gigabytes of memory. However, scDesign2 takes much longer than other methods, and SymSim requires much more memory than the others. This puts an emphasis on the balance between the accuracy of the model and the computational efficiency. scDesign2, for example, explicitly captures the gene-gene correlation, but at the cost of runtime, spending more than 10 hours to simulate 1,344 cells. In contrast, Splatter and SPAR-Sim take a much shorter time to simulate, and their runtime does not differ with the size of the reference data, which demonstrates that sampling each feature from statistical distribution is quicker, but sacrifices simulation accuracy. In general, GLMsim is in the middle tier among the simulators in terms of computational time, and its runtime is stable, the curve not changing much with the complexity of the dataset (Fig. 5.7a). Even the most complex CLL dataset, with multiple cell types, batches and their associations, does not require more time for GLMsim. In addition, GLMsim has the lowest memory usage, especially for the non-UMI dataset.

**Fig. 7.**
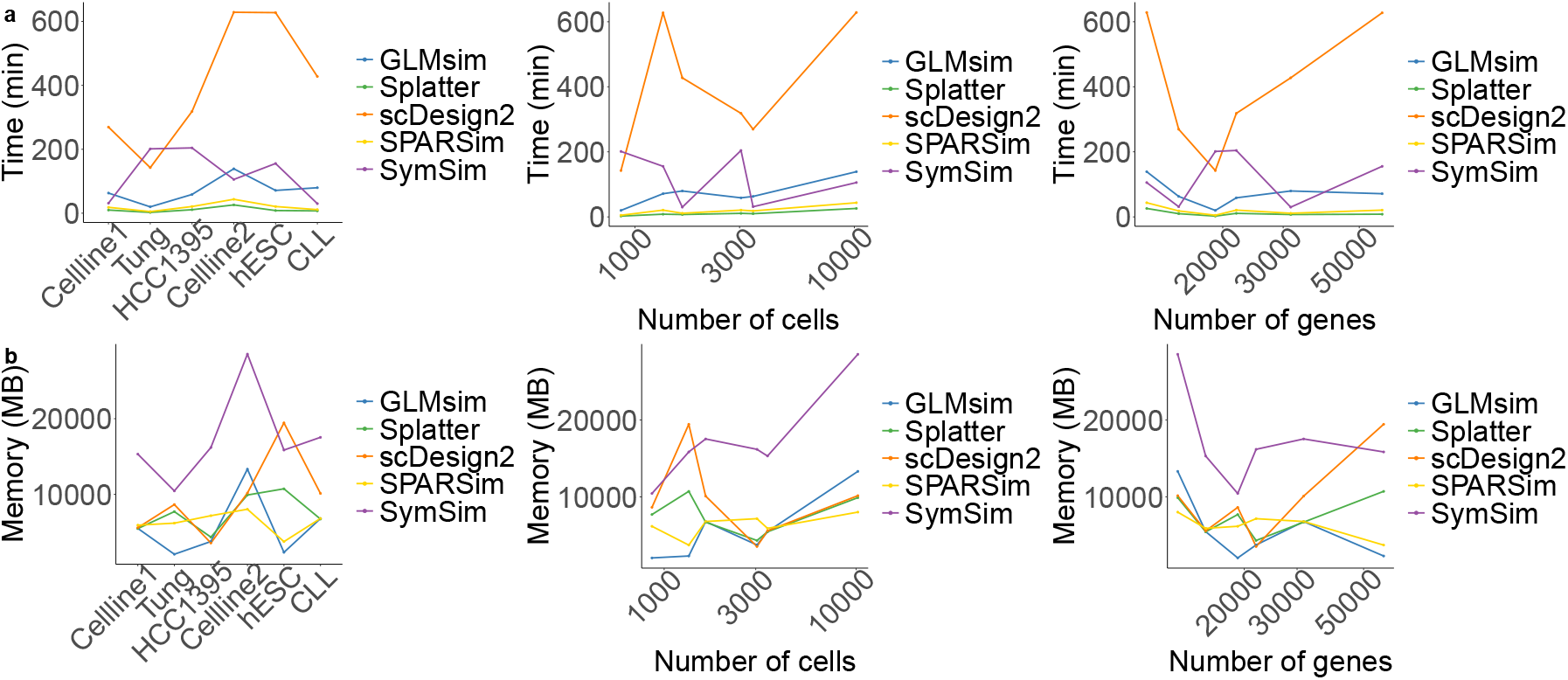
Computational scalability of different simulation methods. (a) Runtime of different methods across all datasets. (b) Memory usage of different methods across all datasets. (a,b) The scalability is also measured by the scale of the data: the number of genes and cells.

### GLMsim is robust to outliers

Some datasets will give extreme simulation values if outliers are not dealt with properly. For example, there is one extreme outlier gene after an initial simulation by GLMsim (Fig. 8a), which is caused by poorly estimated parameters from the Celline1 data. The estimated parameters intercept 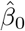, biological coefficient 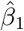, library size coefficient 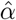 and the dispersion 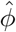 are -9.43, 28.88, 0.66 and 2.18 × 10^*−*4^ respectively. All estimated parameters are within a reasonable range except for 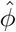 (Fig. 8e), and that leads to an abnormally large negative binomial variance (Equation 3). Consequently, the simulated negative binomial counts can be unreasonably large integers that shift the mean gene expression out of a realistic range.

**Fig. 8.**
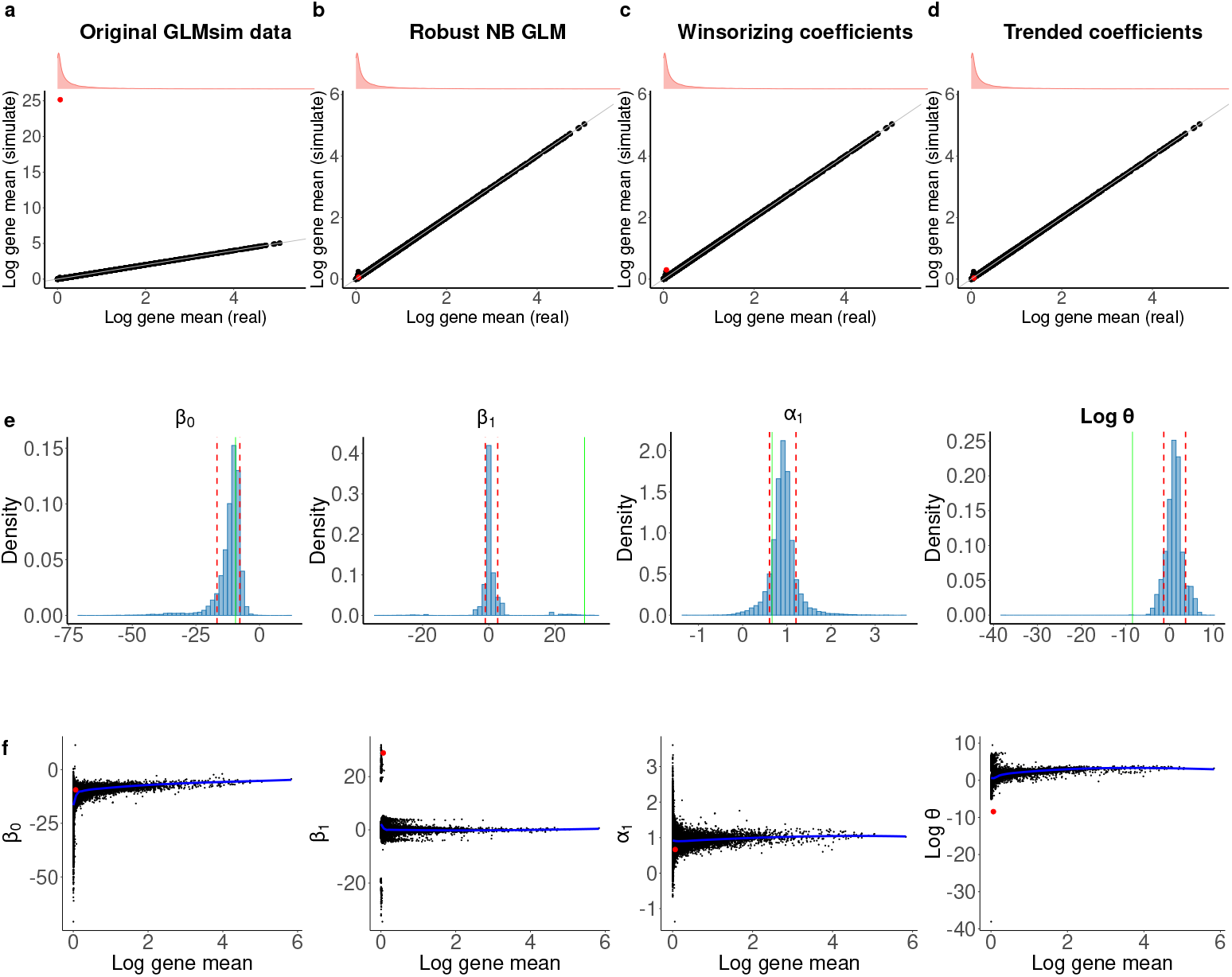
GLMsim handles outlier genes in the Celline1 dataset. The outlier gene is shown by the red dot. (a-d) The comparison of log mean expression between original data and GLMsim simulated data. (a) GLMsim simulated data is from the original simulated data without handling the outlier. (b) GLMsim simulated data after robust NB GLM dealing with outlier gene. (c) GLMsim simulated data after winsorizing the coefficients for the outlier gene. (d) GLMsim simulated data by the trended coefficients to the outlier gene. (e) The distribution of each estimated coefficient across genes. The red lines are cut-offs for each estimated coefficient, which is Q(0.1) and Q(0.9) for each coefficient. The green line is the estimate coefficient for the outlier gene. (f) The relationship between each estimated coefficient and the log gene mean. The blue line represents the loess trended line.

GLMsim provides three strategies (see “Methods”) to address outlier issues. The first method is based on a robust negative binomial GLM, which utilizes the estimated coefficients from robust negative binomial as starting values to refit negative binomial GLM. After applying this strategy, the gene mean expression was assigned to an acceptable level (Fig. 8b). The second approach is winsorizing the fitted coefficients. We found that the *β*_1_ coefficient and the dispersion parameter *ϕ* are beyond the thresholds (Fig. 8e). Then we use the thresholds directly as the new estimated coefficients, which are Q(0.9)(2.68) and Q(0.1)(0.27) for the distributions of *β*_1_ and log *ϕ* across genes, respectively. Finally, the clipped coefficients give counts with a sensible gene mean (Fig. 8c). The third strategy is to construct a relationship between gene means and estimated coefficients in the reference data by fitting a loess line. Now, each predicted coefficient is from the loess line corresponding to the mean expression of the outlier gene (Fig. 8f). Eventually, the outlier gene mean value is optimized when correcting the outlier values by each loess trend across genes (Fig. 8d). In addition to the gene mean, we also examine the library size before and after rescuing the outlier genes (Fig. S2). The simulated library sizes are closer to those from reference data after refining the outliers.

The outlier problem also exists in two other datasets: hESC and CLL. We applied all three methods to those datasets and found that the robust negative binomial GLM and trended coefficient strategies performed stably compared to the winsorizing strategy (Fig. S3a,b). The total computational time among the three methods showed no differences (Fig. S4). Considering the robustness and stability of the three methods, we chose the trended coefficient as the default outlier handling strategy.

### Assessing single cell integration methods using GLMsim

In the original Celline2 dataset, the cells are separated into the Jurkat and 293T cell types, and each cell type is included in two different batches, one alone and another together, which creates an association between biology and batch. We used our simulated Celline2 dataset to evaluate the performance of different scRNA-seq integration methods. A good integration method should remove all the unwanted variation across the batches but keep all the biology. We use different metrics (Supplementary Methods) to assess the extent to which the integration methods achieve these goals.

In the GLMsim simulation of the dataset, the cells are separated into two different biological groups and three different batches along the first principal component (Fig. 9). Using plots of PC2 versus PC1 of the integrated simulated data, RUV-III-NB[26] and scMerge[27] are seen to successfully remove the batch differences and keep the biology. Other methods, such as scran[29], mnnCorrect[24], fastMNN[25], Seurat[30] Pearson residuals and Seurat log corrected data, exhibit no differences before and after data integration, as the batch differences remain. ZINB-WaVE[31] and Seurat Integrated data overcorrect in removing the batch effects, as biology has been removed. In summary, RUV-III-NB and scMerge maintain the biology even when it is associated with batches.

**Fig. 9.**
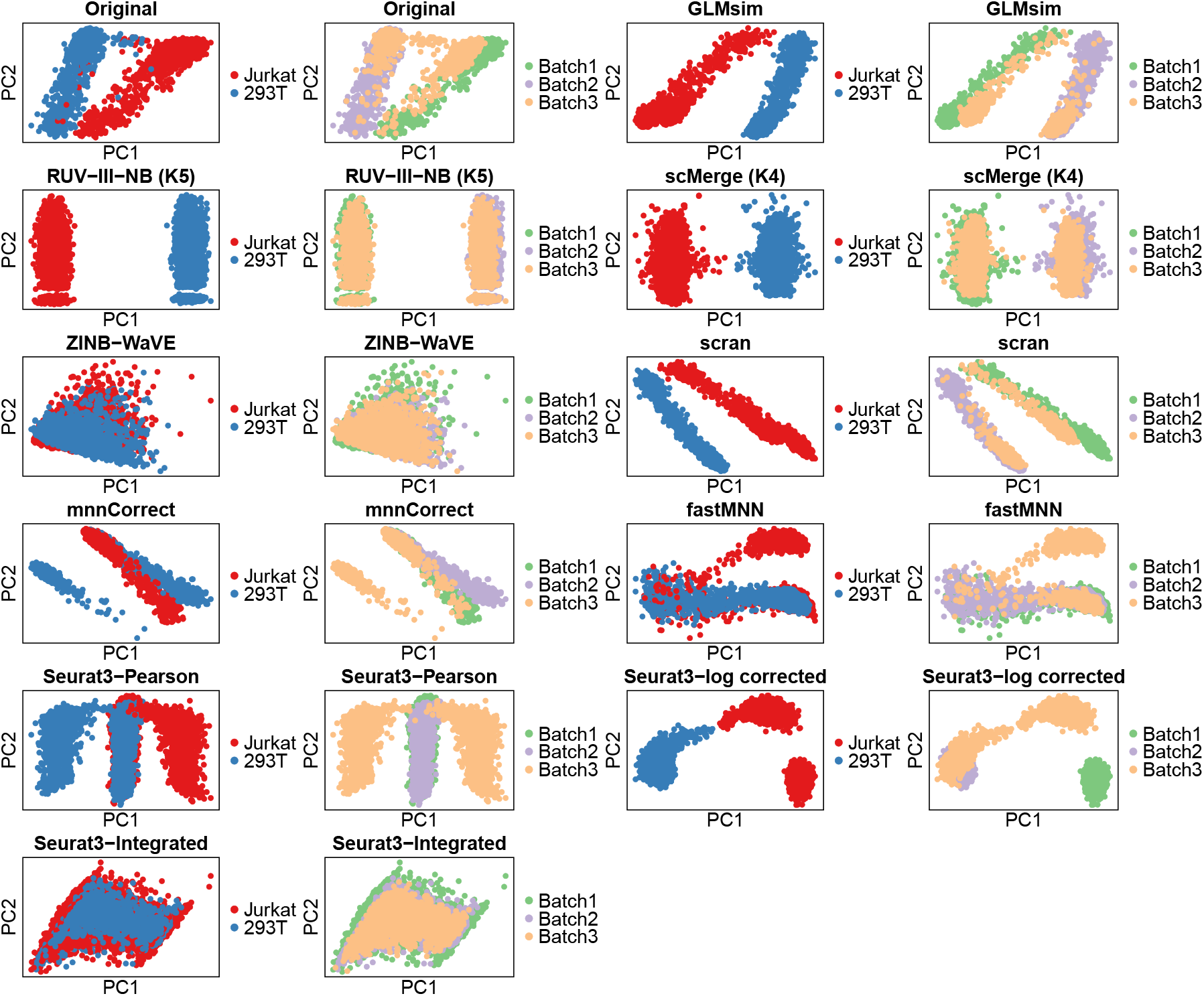
PCA plot of integration on simulated Celline2 data by different methods. GLMsim was used to simulate the Celline2 dataset, then different integration methods were used on the simulated data. The first two principal components are shown in the plot. The cells were colored by cell lines and batches. The first pairs of plots show the original GLMsim simulated Celline2 data. For RUV-III-NB, the plot is made based on log PAC, the log of the percentile-adjusted counts. For Seurat, three data formats are used to do the plots: the Pearson residual, log corrected data and integrated data.

The biological silhouette score, a score ranging from -1 to 1 indicating whether biological groups are clearly distinguished from each other or not, was used to evaluate the ability to enhance biological patterns (Fig. S5). We identified that the RUV-III-NB and scMerge integrated data show their abilities to effectively detect the biological signals, while ZINB-WaVE, fastMNN, mnnCorrect, Seurat Pearson residual and Seurat integrated data fail to pick up the biological signals after data integration. The Seurat log corrected data and scran also performed well to keep the different biological groups. The relative log expression (RLE) plot[32] was then applied to evaluate the performance of removing library sizes. RUV-III-NB and scMerge are the top ranked methods for the RLE metrics (Fig. S6). Except for those two methods, scran also performed well to remove the library size effect although this method is straightforward to reduce the effect by scaling the library size directly. Other methods, such as Seurat, ZINB-WaVE, fastMNN and mnnCorrect still have high correlation with library size after batch removal, which indicates that those methods have limitations in mitigating the library size effect. Another metric to determine the performance of removing library size effects is the Pearson correlation between library size and gene UMI counts. scMerge is the best method to remove the library size effects, because its range of correlation is narrower than other methods (Fig. S7). Other methods performed similarly with correlations close to 0 for almost all genes. fastMNN only provides a data format in low dimensions for visualization purpose. As a result, it cannot be used for down-stream analysis and performed badly with a wider range of correlation to library size. In regard to the batch effects, the proportion of DE genes across batches has been applied to benchmark different integration methods. Theoretically, the proportion of DE genes for the same cell type should be low after removing the batch effects. RUVIII-NB is the unique method that shows low proportion of DE genes, suggesting that RUV-III-NB log PAC data is an ideal choice to carry out downstream DE analysis (Fig. S8). All other methods exhibit a high proportion of DE genes after integration data, indicating that those methods are unable to provide appropriate integrated data format used for downstream analysis.

Overall, RUV-III-NB outperforms other methods in gaining high scores across all metrics (Fig. S9), which implies that RUV-III-NB successfully removed the library size effect, the batch effect and retained the biological effect in the simulated data. In contrast, other methods obtained a low score for at least one metric. For example, the Seurat integrated data is another example that performed badly in almost all metrics except for the technical silhouette score. It illustrates that the Seurat integrated data has the advantage of removing library size in the principal components, but no more benefits are shown by this method.

### GLMsim exhibits simulation stability

It is possible that the random numbers generated by a single cell simulator can influence the simulation results and will further influence the downstream analysis results. In order to check the random effect by the simulator, we simulated 5 Celline2 datasets by setting different random seeds. Then we compared the benchmarking results from the original and all 5 simulated datasets (Fig. S10-S15). We found that the original and all simulated data showed similar performances for all benchmarking metrics. This indicates that GLMsim simulated data is stable and will not be influenced by random aspects of the simulations. Since GLMsim can capture most of the basic features of the original data, the benchmarking results from the simulated data are similar to those from the original data.

## Discussion

In this paper, we have proposed GLMsim, a practical method to simulate the library size, biological and batch effects present in scRNA-seq data. Currently, none of the existing scRNA-seq simulation methods are able to capture associations between these three factors, despite numerous experimental datasets exhibiting such associations. GLMsim achieves this goal by incorporating library size, batch and biology in the model. In this way, GLMsim not only recovers the information relevant to these three factors from experimental data, it also efficiently handles challenging large-scale datasets with multiple batches and biological groups. Since most single cell simulators simulate the three factors separately in their models, the simulation patterns for those methods will be poor representations of the actual data, especially with complex scenarios. In particular, if the method simulates different single cell groups through multiplying by DE factors, the assignment of DE genes to highly diverse batch and biological clusters will be problematic, because those methods cannot avoid the DE assignments across different clusters.

We have compared GLMsim to other single cell simulators by a series of gene-level and cell-level summaries to evaluate the performance of GLMsim in terms of its ability to capture the characteristics of experimental data and its fidelity to that data. Utilizing 6 datasets with different numbers of genes and cells, sequenced by different platforms, we found that GLMsim ranked best in simulating data similar to experimental data. In particular, GLMsim is the only method that enables us to precisely reproduce the cell level proportions of zeros in non-UMI data. scDesign2 and SPARSim performed well in the gene-level metrics, but poorly in simulating cell-level features. Splatter and SymSim have a poor performance in every respect. In summary, GLMsim is the most accurate single cell simulator of basic single cell properties, with the notable exception of gene-gene associations. Its accuracy lies in the GLM being able to estimate parameters for each gene.

Single cell data typically includes 10-20 thousand of genes and at least hundreds, if not thousands, of cells. As a result, it can be time consuming fitting every gene using a GLM and so. We parallelized the computations when estimating the parameters, and we also provide a sequential option for users to obtain the fitted coefficients in case of runtime errors. Although GLMsim is slightly slower than the distribution-based methods such as Splatter and SPARSim, GLMsim successfully balances the accuracy of simulation and the computational time. Moreover, GLMsim has lower memory usage than the other methods, especially for non-UMI data.

Current single cell simulators ignore or filter out outlier simulated values. GLM-sim addresses this problem with three different approaches instead. It offers a robust negative binomial GLM, the winsorizing of estimate and the trended approaches. All these methods address the outlier problem well, but the performance of the trended method is more stable than the other two methods. The runtime of the three strategies does not differ significantly, and thus, we choose the trended coefficient approach as the default option.

GLMsim simulated data has been applied to give comprehensive benchmarking across popular single cell integration methods. We have shown that RUV-III-NB outperformed other methods in most of the metrics, while scMerge was in second place, as it is slightly weaker in removing batch effects on the metric relevant to the proportion of DE markers across batches. Some of the methods do not show any differences before and after integration, such as scran, mnnCorrect, fastMNN, and Seurat, suggesting that those integration methods lack the ability to deal with library size, batch, and associations between batch and biology. On the other hand, other methods demonstrate overcorrection problems, like ZINB-WaVE and Seurat integrated data. We also found that some integration methods can only provide data in reduced dimensions for visualization purpose, including mnnCorrect and fastMNN. However, those integrated data in low dimensions cannot support for certain downstream analyses such as detecting DE markers. In addition, we have demonstrated that GLMsim performs stably on benchmarking, since same conclusions were given across simulations when offering different random seeds to simulate the same real dataset. Benchmarking results based on the simulated data offer researchers an objective standard with which to select an appropriate approach for single cell analysis.

GLMsim simulation is robust, reproducible, user-friendly, and the framework is distinctive compared to any other method. Users only need four steps to finish simulation. First, provide basic information from experimental data, which includes the count matrix and the biological and batch labels for each cell. Second, it estimates parameters for each gene. This step is time-consuming, so we encourage users to carry it out on a high performance computer system if possible. Next, simulate an initial count matrix, which requires the parameter estimates from the previous step to calculate the estimated mean for each entry of the matrix. Finally, check if outlier genes exist in the initial simulated data, and if so choose an appropriate method to correct for these outlier values. At present, GLMsim only works for simulation of scRNA-seq data.

## Methods

### Estimating coefficients for each gene

Let 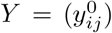 represent the count matrix from the original dataset, whose *G* rows correspond to genes and *N* columns correspond to cells, respectively. Write

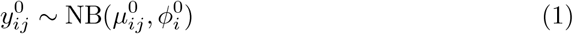

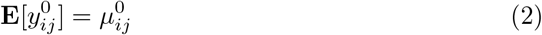

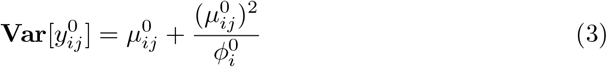

to denote that for gene *i* in cell *j*, 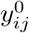 is distributed according to a negative binomial (NB) distribution[33] with mean parameter 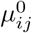 and dispersion parameter 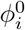.

Assume that in the original data, there are *M* biological groups and *K* batches. Our GLM then takes the form

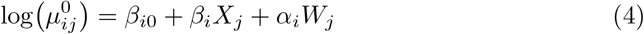

where *β*_*i*0_ is the baseline expression of gene *i* for a reference biological group and reference batch group, *β*_*i*_ is a vector of parameters for the biological influences and *α*_*i*_ is a vector of parameters relevant to the unwanted variations. *X*_*j*_ = (0, *x*_*j*2_, · · ·, *x*_*jM*_)^*T*^ is a vector of parameters related to the biological groups that if cell *j* belongs to the *m*-th group other than the reference group, *x*_*jm*_ = 1 and other entries are 0. *W*_*j*_ = (*L*_*j*_, *w*_*j*1_, · · ·, *w*_*jK*_)^*T*^ is a vector of unwanted variation including library size and batches. *L*_*j*_ corresponds to log library size for cell *j*:

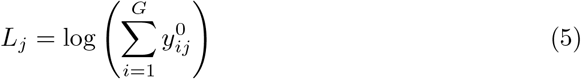

*w*_*jk*_ corresponds to batch for cell in non-reference groups that if cell *j* is in the *k*-th batch, *w*_*jk*_ = 1 and other entries are 0. We use the `glm.nb`[33] function from the package `MASS` to get the estimated parameters: 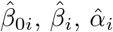 and 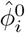 for each gene from the dataset being fitted.

## Simulating single cell gene counts

In this step, the counts for each gene are simulated independently using the estimated coefficients from the previous step. The estimated mean expression parameter 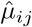 of the simulated count *y*_*ij*_ for gene *i* in cell *j* is defined as:

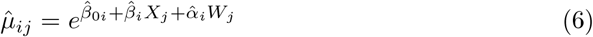

After computing the estimated mean expression for the count to be simulated, we sample the counts from either the negative binomial distribution or the Poisson distribution. The majority of genes are able to get an estimate 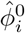 from the data, and their simulated counts, *y*_*ij*_, can then be drawn from the negative binomial distribution:

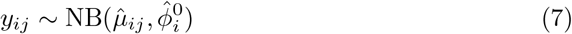

However, a small proportion of genes fail to return an estimated dispersion 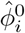, which is likely caused by their dispersion characteristics. In such cases, we use the Poisson distribution to simulate their counts:

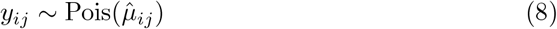

### Rescuing outlier genes

It is possible to introduce outlier values in the initial simulation of gene counts. Thus, in this additional step, we aim to check if outlier gene counts exist. If they do, we can use one of three optional methods to correct them. We check for outliers by comparing the mean gene expression levels of the simulated data to those of the original data. We define the mean expression of gene *i* from the simulated data by *λ*_*i*_ and the original data 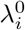 by:

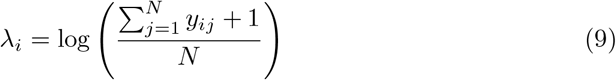

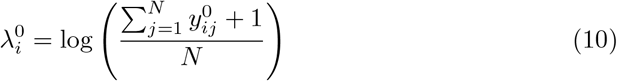

For each gene, we can get the absolute difference 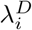 between the simulated mean expression and real expression:

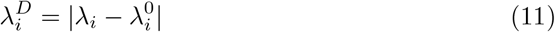

Then we obtain the median absolute deviation *λ*_MAD_ of 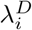 across all genes. The genes with 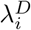 larger than a chosen cut-off are labelled as outliers. That is, gene *i* is an outlier if 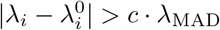. The default value for *c* we use is 30.

#### •Robust negative binomial

The first way to rescue outliers is the robust negative binomial method[34]. Most of outliers are caused by poorly estimated 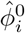. Outlier genes are refitted using a robust negative binomial regression model with the same design matrix as *X*_*j*_ in (4), and again obtain the refitted coefficients 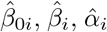, and 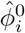. Then we refit a classical negative binomial GLM with the `glm.nb` function but using the above robustly estimated coefficients as the starting values. This gives a new set of estimated coefficients and we use them to update the estimated mean and sample gene counts.

#### •Winsorizing

The second strategy to deal with outliers is winsorizing. For each fitted coefficient, we set a cut-off based on the quantile of its distribution across all genes. The default cut-offs are the 5% quantile Q(0.05) and the 95% quantile Q(0.95) of the distribution. If the coefficient of the outlier gene falls in the top or bottom 5% of the distribution, we use the cut-off value directly as its new fitted coefficients. For example, for an outlier gene *g*, suppose we find that its estimated coefficients 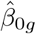 is within the range of 5%-95% of the distribution 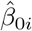 across all genes, and the same for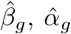, then we keep those three estimated coefficients. But if we find that log 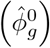 is outside the 5%-95% of the distribution of log 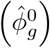, and is closer to the Q(0.05) of that distribution, we set the exponential of Q(0.05) as the new 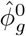. In the subsequent steps, negative binomial counts for gene *g* are sampled using 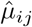 with these new fitted coefficients, giving revised simulated gene counts.

#### •Trended coefficient

The trended coefficient approach is the default method for handling outliers. We construct the relationship across genes between 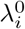 and each gene’s coefficient by loess regression. Notice that here we use a logarithmic transformation for the dispersion parameter 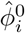. For outlier genes, the loess smoothed value are their new estimates. After that, counts are drawn from negative binomial distribution with the estimated 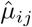 computed using the corrected parameter estimates.

### Benchmarking different single cell simulation methods

The version of Splatter is 1.20.0, and the version of scDesign2 is 0.1.0. We also used version 0.9.5 of SPARSim and version of 0.0.0.9000 of SymSim for benchmarking purpose. The definitions of the features compared follow. We denote the raw simulated count matrix by *Y*_*G×N*_, where *i* refers to gene *i* in rows and *j* refers to cell *j* in columns. Assume there are *G* genes and *N* cells in this count matrix. Then the log gene mean *λ*_*i*_ is defined as 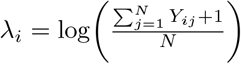. The log library size *L*_*j*_ is: 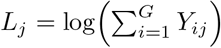. The log gene UMI total is defined as: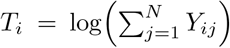. The gene variance is: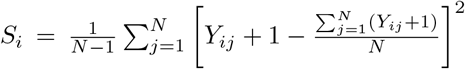. Denote the gene proportions of zeros by 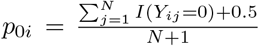. The logit transformation of *p*_0*i*_ is: 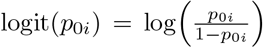 Denote the cell proportions of zeros by 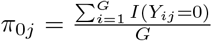. The logit transformation is logit 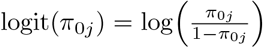. Here, *I*(*Y*_*ij*_ = 0) = 1 if *Y*_*ij*_ = 0 is true and *I*(*Y*_*ij*_ = 0) = 0 otherwise.

### Datasets

In order to benchmark different single cell simulators, we use six datasets sequenced by different platforms and include different scenarios for biological groups as well as batches (Table S1). All datasets start from the raw single cell count matrix without pre-processing. For the hESC dataset some extremely low abundance genes may inappropriately bias the estimation of the GLM parameters, hence the genes expressed in less than 4 cells were filtered out of this dataset. For the other datasets, we use the raw scRNA-seq counts directly.

#### Dataset 1: Celline1

The cells in the dataset[35] are a 50-50 mixture of Jurkat and 293T cells in one batch. This dataset is batch 3 of the Celline2 dataset. It is used to study a particular biological issue.

#### Dataset 2: Tung

The Tung dataset[36] was generated on the Fluidigm C1 platform and was used to explore the sources of technical variation in scRNA-seq technology. The data was collected from induced pluripotent stem cells (iPSC) of three Yoruba samples (NA19098, NA19101, NA19239). Each sample was independently collected three times, and each replicate was processed using the same reagents. ERCC spike-in controls were added to each sample. The samples were sequenced by the SMARTer protocol. The data is available via: https://github.com/jdblischak/singleCellSeq

#### Dataset 3: HCC1395

The 10x breast cancer cell line dataset[37] was used as a benchmarking dataset to compare different single cell methods. The cells were collected from a 43-year old female donor. We selected the pure HCC1305 cells sequenced at Loma Linda University. The cells in this dataset have a wide range of library sizes and are used for exploring the library size-only effects. The dataset was downloaded from: https://springernature.figshare.com/collections/A_Multi-center_Cross-platform_Single-cell_RNA_Sequencing_Reference_Dataset/5213468

#### Dataset 4: Celline2

The dataset[35] was produced for the purpose of investigation of the 10x platform. The cells come from two quite different cell lines: Jurkat and 293T. There are three batches in the dataset. One batch is all Jurkat cells; another batch is all 293T cells, and the third batch is a 50:50 mixture of Jurkat and 293T cells. The three batches were pre-processed separately using the same standard, which involved preserving the features expressed in at least 10 cells and detecting at least 200 genes in each cell.

The batch1 count matrix was downloaded from: https://www.10xgenomics.com/welcome?closeUrl=%2Fresources%2Fdatasets&lastTouchOfferName=Jurkat%20Cells&lastTouchOfferType=Dataset&product=chromium&redirectUrl=%2Fresources%2Fdatasets%2Fjurkat-cells-1-standard-1-1-0

The batch2 count matrix was downloaded from: https://www.10xgenomics.com/welcome?closeUrl=%2Fresources%2Fdatasets&lastTouchOfferName=293T%20Cells&lastTouchOfferType=Dataset&product=chromium&redirectUrl=%2Fresources%2Fdatasets%2F293-t-cells-1-standard-1-1-0

The batch3 count matrix was downloaded from: https://www.10xgenomics.com/welcome?closeUrl=%2Fresources%2Fdatasets&lastTouchOfferName=50%25%3A50%25%20Jurkat%3A293T%20Cell%20Mixture&lastTouchOfferType=Dataset&product=chromium&redirectUrl=%2Fresources%2Fdatasets%2F50-percent-50-percent-jurkat-293-t-cell-mixture-1-standard-1-1-0

#### Dataset 5: hESC

Naïve and primed human embryonic stem cells (hESCs) were profiled to investigate the heterogeneity and developmental transition within each pluripotency state[38]. Naïve hESCs were grown in N2B27 medium with titrated 2 inhibitors (PD0325091 and CHIR99021), Leukemia inhibitor and Go6083 inhibitor, while primed hESCs were cultured in E8 media. Naïve hESCs were processed in two batches: the first batch contained 96 cells in each state, and the second batch contained 384 cells in each state. Primed hESCs are in the same condition for the first two batches as the naïve hESCs, but the primed cells have an additional 384 cells in a third batch. The cells were prepared and sequenced by the SmartSeq2 protocol. The data is available at: https://bioconductor.org/packages/release/data/experiment/vignettes/scRNAseq/inst/doc/scRNAseq.html

#### Dataset 6: CLL

This dataset[39] was part of an investigation of Venetoclax (VEN) resistance. The majority of cells are B cells. The cells were treated with dimethyl sulfoxide (DMSO), VEN and combinations of VEN and other treatments for one week. The data was generated on the CEL-Seq2 platform over 6 plates (LC89, L91, L93, L95, L96, LC99). Granta cell line cells were included in each plate. This dataset is the most challenging one for three reasons. Firstly, it has multiple batches and biological groups. Secondly, associations exist between library size, batch and biology. Lastly, except for the Granta cell line, each drug treatment condition is dominant in one batch. In other words, different cell types are not evenly mixed in each batch. This dataset can be accessed by requesting permission from the authors.

## Declarations

### Ethics approval and consent to participate

Not applicable

### Consent for publication

Not applicable

### Availability of data and materials

See the section of “Dataset”. The code is available at Github (https://github.com/jiananwehi/GLMsim.git). The R package is under a GPL-3.0 license.

### Competing interests

The authors declare that they have no competing interests.

## Funding

Jianan Wang is supported by Walter and Eliza Hall International Scholarship and CSL Translational Data Science Scholarship.

## Authors’ contributions

JW developed the method and conducted all analysis. LC contributed to statistical methods for GLM parameter fitting and non-UMI data. RT designed and generated data for the CLL study. BP contributed to trended coefficients and supervision. TS oversaw the whole project. All authors read and approved the final manuscript.

## Acknowledgements

We would like to thank to Givanna Putri for providing suggestions of parallel computation. Our thanks also to Chris Woodruff and Luke Gandolfo for their comments on the manuscript.

## Notes

### Competing Interest Statement

The authors have declared no competing interest.

